# Brain structural covariances in the ageing brain in the UK Biobank

**DOI:** 10.1101/2022.07.26.501635

**Authors:** Chao Dong, Anbupalam Thalamuthu, Jiyang Jiang, Karen A. Mather, Perminder S. Sachdev, Wei Wen

**Affiliations:** Centre for Healthy Brain Ageing (CHeBA), Discipline of Psychiatry and Mental Health, UNSW Medicine & Health, University of New South Wales, Sydney, New South Wales, 2052, Australia; Neuroscience Research Australia (NeuRA), Sydney, New South Wales, Australia; Neuropsychiatric Institute (NPI), Prince of Wales Hospital, Randwick, New South Wales, 2031, Australia

**Keywords:** Brain ageing, structural MRI, structural covariance, cognition, longevity-PRS

## Abstract

Brain structural covariances or pairwise correlations describe how morphologic properties of brain regions are related to one another across individuals. Although it is reported that brain structural covariance changes during brain development, it is not clear how structural covariance relates to the ageing process. Here we investigated the human brain structural covariances of cortical thickness and subcortical volumes in the ageing brain and their associations with age, cognition, and longevity polygenic risk score (longevity-PRS) by using cross-sectional data from the UK Biobank (N = 42075, aged 45-83 years, M/F=19752 /22323). The sample of participants was divided into 84 non-overlapping groups based on their age. The older the age group, the greater the variability in the whole brain structural covariance. However, there was a differential rate of age-related increase of variance between males and females. The variance of females started lower than those of males and then increased with age with a greater gradient than that of males. There was a consistent and significant enrichment of pairwise correlations within the occipital lobe in ageing process. The cortical thickness and subcortical covariances in older groups were significantly different from those in the youngest group. Sixty-two of the total 528 pairs of cortical thickness correlations and 10 of the total 21 pairs of subcortical volume correlations were significantly associated with age after Bonferroni correction. Specifically, with an increasing age, most decreased cortical thickness correlations were found between the regions within the frontal lobe as well as between the frontal lobe regions and regions in other lobes, while pairwise correlations within occipital lobe regions were all strengthening. Most of these correlations were also associated with global cognition and weakly associated with longevity-PRS. These findings revealed that the structural covariance was not stable during ageing. Given the thinning of the cortex and the volumetric reduction of subcortical structures seen in the ageing process, an increased pairwise correlation between the brain regions in the older brain suggested a strengthened coordinated decline between the brain regions involved. However, some of the brain regions demonstrated a differentiated rate of decline which was shown as the inversed or reduced pairwise correlations between these regions.

## 1. Introduction

Morphologic properties of brain regions covary with each other, which can be modelled by brain structural covariance (Mechelli et al., 2005). It has been increasingly recognised that brain structural covariance reflects a synchronised maturation process in the developmental stages of early childhood and adolescence brains and a coordinated atrophy and decline in the ageing brain. However, the exact biological meaning of this structural covariance still remains controversial (Alexander-Bloch et al., 2013). Magnetic Resonance Imaging (MRI) is a non-invasive, widely used technique that enables the observation of living human brain anatomy (Lerch et al., 2017). Macroscale structural covariance can be estimated by T1-weighted MRI data, from which brain morphology measures such as cortical thickness and volume, are obtained (Lerch et al., 2017). These morphological covariances are associated with axonal and synaptic connectivity between brain regions, with shared genetic and neurodevelopmental effects (Alexander-Bloch et al., 2013; Grasby et al., 2020; Hofer et al., 2020; Strike et al., 2018). To date, however, there is still a lag in understanding the specific changing patterns of structural covariance in the ageing brain, and how they relate to cognition and longevity.

The simplest way of defining structural covariance is to estimate the correlation between the morphological property of two brain regions at the group-level (Carmon et al., 2020). The ‘structural covariance matrix’ means the correlation matrix of the whole brain (Carmon et al., 2020). Instead of analysing the regional pairwise covariance directly, many studies have been exploring brain covariance-based network measures (Bullmore and Sporns, 2009, 2012; Evans, 2013). Age-related changes in structural covariance may be reflective of a disturbance of the biology of brain morphologies. Investigating brain covariance principles can advance our understanding of the patterning of brain structures and help investigate relevant diseases such as Alzheimer’s disease (Montembeault et al., 2016; Nestor et al., 2017). For example, a previous study explored age-related differences in grey matter density structural covariance in the different developmental stages of childhood and adolescence by dividing the participants into four equally sized age groups (Zielinski et al., 2010). Similarly, other studies found that cortical thickness covariance also had age-related changes during adolescence by applying sliding-window configurations, suggesting a mixture of linear and non-linear alterations during the period of maturation (Váša et al., 2017; Vijayakumar et al., 2021). However, so far, most studies about age-related structural covariance in the human brain have focused on the development of the brain during childhood and adolescence (Sotiras et al., 2017), leaving much to be learned for the ageing process. Additionally, the sample sizes in these studies were generally not large enough.

It has been found that changes of cortical thickness covariances in both development and ageing were closely correlated with the genetic organisation (Fjell et al., 2015), and subcortical structures were also governed by the same sets of genes (Fjell et al., 2021). It would be interesting to explore whether cortical thickness and subcortical covariances are associated with longevity by examining relevant genes of longevity polygenic risk scores (longevity-PRS) (Dong et al., 2022; Tesi et al., 2021). Additionally, as cognition usually declines in the ageing process (Bishop et al., 2010; Grady, 2012) it is therefore important to understand the relationship between the brain covariance and cognition.

In this study, we aimed to investigate brain structural covariances, including cortical thickness and subcortical volume covariances in the ageing process and explore the associations between these covariances with age, cognition and longevity-PRS. As structural covariance needs to be estimated at group level, we divided our sample into age-specific groups (~500 participants in each group). Within each group, we then estimated the covariance in cortical thickness and in subcortical volumes separately. We hypothesised that structural covariance would show age-related properties and would be also associated with cognition and longevity-PRS.

## 2. Methods

### 2.1 Participants

All our data were drawn from the UK Biobank, which is a population-based study consisting of over 500,000 participants aged between 40-70 years at study entry (Sudlow et al., 2015). Written consent was acquired from all participants and ethics approval was provided by the National Health Service National Research Ethics Service (11/NW/0382). Imaging data and cognition data in the current study were acquired at the first imaging assessment visit (instance 2). Here we used the MRI data released in 2021 under the UK Biobank application number 37103. After removing outliers, i.e., data points greater than five standard deviations from the mean, 42075 participants were included in the final analysis. The age range of the participants was 45 to 83 years with the mean of 64.5 years (Table 1).

### 2.2 Neuroimaging measures

Brain anatomical measures of cortical thickness and subcortical volumes were derived from structural MRI scans (Siemens Skyra 3T with a standard Siemens 32-channel RF receive head coil). The sequence parameters of T1-weighted structural imaging were: spatial resolution 1×1×1mm; field of view 208×256×256 matrix; TI (inversion time) 880 ms; TR (repetition time) 2000 ms. The full protocol is available at http://biobank.ctsu.ox.ac.uk/crystal/refer.cgi?id=2367. We used data that were preprocessed by UK Biobank. Brain cortical parcellation was performed using the Desikan-Killiany atlas (Desikan et al., 2006). We used 33 cortical thickness and seven subcortical volumetric measures for each hemisphere.

### 2.3 Cognition

Cognitive assessments were administered on a fully-automated touchscreen questionnaire (Sudlow et al., 2015). Seven tests from the UK Biobank battery of cognitive tests were selected for the current study to represent three cognitive domains: processing speed, executive function, and memory (Cox et al., 2019; Kendall et al., 2017). All test scores were z-transformed and then averaged to form domain scores. Global cognition was estimated by averaging the domain scores and z-transform. There were 27140 participants with both MRI scans and full cognition data.

### 2.4 Genotyping and Imputation and longevity polygenic risk scoring

Genotyped and imputed data for approximately 500,000 samples were available. The details of genotyping and imputation in UK biobank samples can be found elsewhere (Bycroft et al., 2018; McCarthy et al., 2016). Briefly, the genotyping was performed using the Bileve or UK Biobank axiom arrays. Imputation was undertaken using the Haplotype Reference Consortium (HRC), 1000 genome and UK Biobank reference panels. For PRS calculation only those Single nucleotide polymorphisms (SNPs) which remained after quality control (QC) filtering (MAF > 0.1%, imputation information score > 0.6) were used. In addition, we used only those samples who were reported as British ancestry and removed any samples with high genotype missing rate and relatedness.

Longevity GWAS (Deelen et al., 2019) summary statistics were downloaded from the GWAS Catalog (https://www.ebi.ac.uk/gwas/downloads/summary-statistics). The longevity PRS was calculated using the program PRS-CS (Ge et al., 2019), which uses the Bayesian regression framework with continuous shrinkage priors to obtain posterior effect sizes of the summary statistics. In this study we have used the precompiled disequilibrium (LD) pattern of the 1000 Genome European reference panel with the default parameters of PRS-CS. The pruned set of effect sizes from the whole genome (without the need to use thresholds based on the GWAS p-values) was used to generate PRS scores using PLINK software (Chang et al., 2015). There were 40978 participants who had both MRI data and longevity-PRS data.

### 2.5 Statistical analyses

#### 2.5.1 Construction of structural covariance in each age-specific group

Regional thickness and subcortical volume were defined as the mean of the left and right hemisphere. For the analysis of covariance matrices across age groups:

1. All 42075 participants were sorted according to age from the youngest to the oldest, and were then split into nonoverlapping, equal-sized 84 groups with 500 participants in each group (the last age group included 575 participants).
2. We obtained residuals of regional cortical thickness estimates after regressing out sex, scanner, and global mean cortical thickness, and then calculated within each group pairwise Pearson correlations across all the 33 regions of interest (ROIs), generating a 33×33 correlation matrix map for each group which contained (33×32)/2=528 pairwise correlations.
3. Similarly, residuals for the subcortical volume estimates were obtained after regressing out sex, scanner, and intracranial volume (ICV), and a 7×7 Pearson correlation matrix in each group [(7×6)/2=21 pairwise correlations] was then generated.
4. To investigate if the structural covariance was different in male and female participants, we also estimated cortical thickness and subcortical volume covariances for the same 84 groups in males and females separately without regressing out sex when computing residuals.

All cortical and subcortical regions can be found in Table S1 and Table S2.

#### 2.5.2 Whole brain variability properties of structural covariance across all age groups

To examine the age-related changes of the overall correlation matrices, we have computed three whole brain variability measures of the age group correlation matrix: variance, von Neumann entropy, and proportion of pairs of correlations that differ significantly within the group. The sample variance of whole pairwise correlations and the entropy captured the variability of the correlation coefficients. The proportion of significant pairwise differences within the correlation matrix provided another estimate of variability of the correlation coefficients. These variability measures for each of age group correlation matrices were examined against median age.

Let R denote the Pearson correlation matrix of the cortical thickness (33×33) or subcortical volumes (7×7). The calculation of these three measures were described as following.

##### Variance measure

the sample variance of the lower diagonal elements of the R matrix.

##### von Neumann entropy

*S*(*ρ*) = –*Trace*(*ρ* log(*ρ*)) =–∑*λ_j_* log (*λ_j_*), 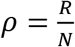 where N is the number of ROIs in the correlation matrix (N=33 or 7) and *λ_j_* are the eigen values of the *ρ* matrix (Felippe et al., 2021). The entropy takes minimum value when all the correlation coefficients are unity and reaches the maximum value (log N) if the all the correlations are zero.

##### Proportion

To examine the pair-wise correlation differences with each group, a test for equality of pairs of correlations was performed. The proportions of pairs that differ significantly out of the total number of comparisons (total number of tests=528×527/2= 139,128 for cortical regions and 21×20/2=210 for subcortical regions) were computed from the correlation matrix of each age group. Tests for equality of two dependent correlations measured on the same set of individuals were performed using the Steiger’s method (Steiger, 1980) as implemented in the R package *cocor* (Diedenhofen and Musch, 2015).

In order to investigate if there are sex differences in these 3 measures (variance measure, von Neumann entropy and the proportion of pairs that differ significantly), the statistical differences between males and females about these 3 measures were estimated using an independent samples t-test.

#### 2.5.3 Over-representation of correlation analysis (ORCA)

To test the enrichment of correlation coefficients between and within lobar regions, ORCA (Pomyen et al., 2015) was performed. Four lobes were included here: frontal, temporal, parietal, and occipital (Table S1). Significant enrichment of correlation coefficients above a threshold than by chance between/within lobe regions was assessed using hypergeometric probability distribution. Correlation threshold was determined using the maximum Shannon entropy measure (Shannon, 1948). Enrichment of correlation using ORCA was performed within each lobe and between all possible pairs of lobes (4×3/2=6) for the correlation matrix of each age group. Bonferroni correction was used to adjust for multiple hypotheses tests (n=10; 4 within lobe + 6 all possible pairs of lobes).

#### 2.5.4 Test for equality of correlation coefficients between the youngest age group and each of other groups

Global tests for equality of elements between two independent correlation matrices [the youngest age group (group 1) and each of other groups (group 2-group 84)] were done using the “pattern hypothesis” approach (Steiger, 1980) as implemented in the R application MML-WBCORR (Fouladi, 2018). The test of equality of elements of correlations in the first age group was compared with corresponding elements of the correlation matrices of the other age groups. Similarly, the elements of correlation matrices obtained from using male and female samples separately at each age group was compared.

#### 2.5.5 Associations between structural covariance, age, cognition and longevity-PRS

To test the associations between pairwise correlations and age, we estimated the associations between each element of the correlation matrix (i.e., the upper or lower triangle of matrix) and age (we used median age in each group) across all 84 groups by using Pearson correlation. Specifically, we firstly applied Fisher’s r-to-z transformation to all elements of the correlation matrix and then tested the association between each Fisher’s z-transformed correlation and age across all 84 correlation matrices. Therefore, (33×32)/2=528 associations of cortical thickness correlations and (7×6)/2=21 associations of subcortical volume correlations were estimated. Similarly, we tested the associations between Fisher’s z-transformed correlation and median cognition (processing speed, executive function, memory, and global cognition) and median longevity-PRS across all 84 groups. As some participants did not have cognition or longevity-PRS data, the median cognition and median longevity-PRS were estimated for each group using only the participants with available data. Sex and scanner covariates for this analysis were not used because the correlation matrices were generated after removing the effects of these covariates. For all analyses, a Bonferroni-corrected p-value (cortical thickness covariance: p<0.05/528 = 9.47e-05; subcortical covariance: p<0.05/21 = 0.0024) was considered statistically significant. Statistical analyses were performed using R version 4.1.0.

## 3. Results

### 3.1 Sample characteristics and experimental design

Sample characteristics are shown in Table 1 and the conceptual overview of the study is depicted in Figure 1. Firstly, the structural covariance (correlation) matrices were established in each group. The cortical structural covariance of the first two youngest groups (group 1, 2) and last two oldest groups (group 83, 84) can be found in Figure S1. At the whole brain level, brain variability measures of the age group correlation matrix were examined. ORCA was applied between lobar regions within each of the correlation matrix. We also performed the test for equality of correlation coefficients between the youngest age group and each of other age groups. At the ROI level, associations between each element of correlation matrix, age, cognition, and longevity-PRS across 84 groups were investigated.

**Table 1.** Sample characteristics in UK Biobank (instance 2).

| **Participants with MRI data** |  |  |
| --- | --- | --- |
| | Size | Mean age ± SD (age range) |
| All participants | N = 42075 | 64.5 ± 7.7 (44.6-82.8) |
| Male | N = 19752 (46.9%) | 65.2 ± 7.8 (44.6-82.5) |
| Female | N = 22323 (53.1%) | 63.8 ± 7.5 (45.2-82.8) |
| **Participants with both MRI and cognition data** |  |  |
| All participants | N = 27140 | 64.7 ± 7.6 (47.0-82.8) |
| **Participants with both MRI and longevity-PRS data** |  |  |
| All participants | N = 40978 | 64.5 ± 7.7 (44.6-82.8) |

Instance 2 means the first imaging assessment. SD: standard deviation. Longevity-PRS: longevity polygenic risk score.

**Figure 1.**
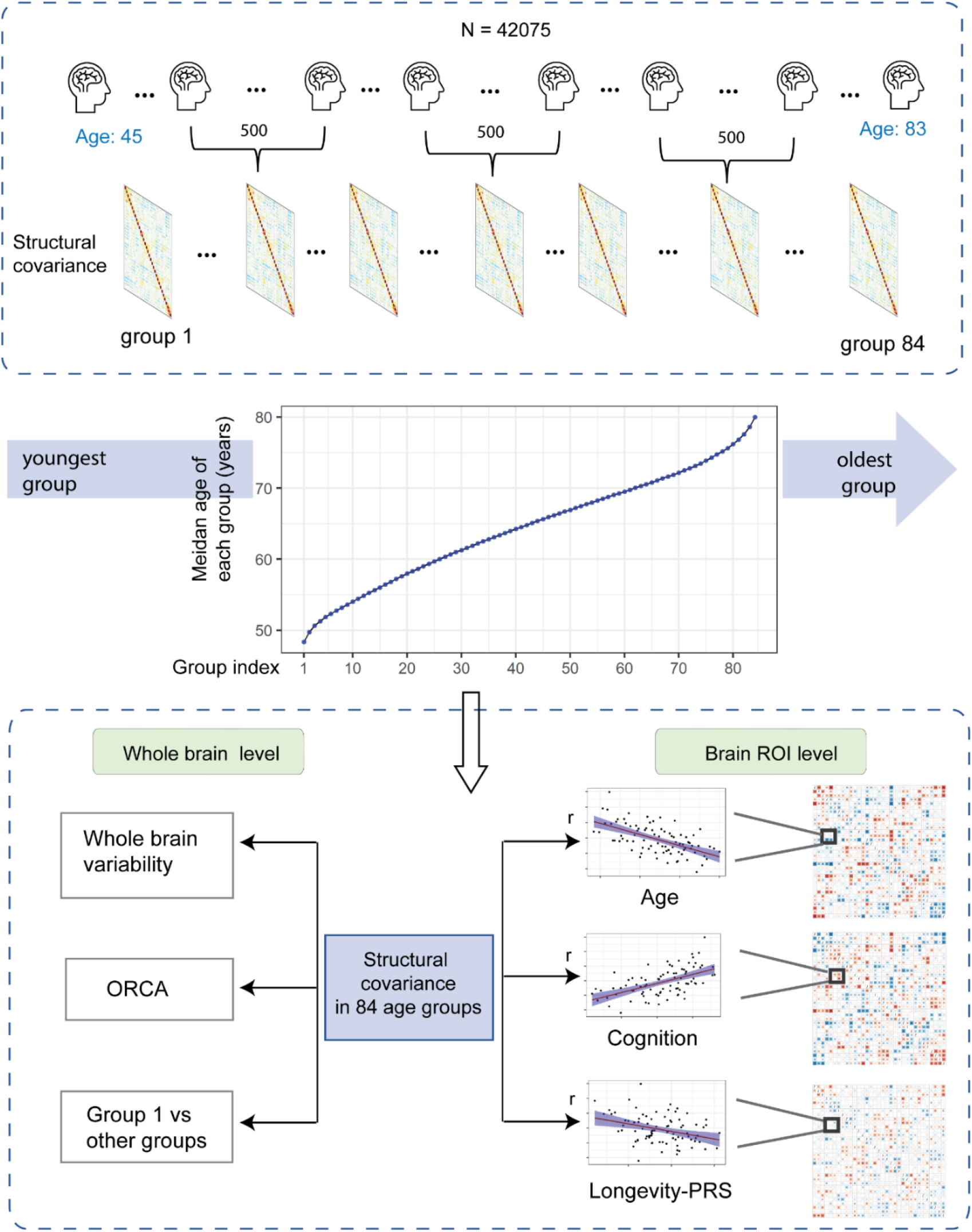
Conceptual overview of the study. Firstly, all 42075 participants were ordered from the youngest to the oldest. They were then divided into equally sized 84 groups with 500 participants in each group. Structural covariance was estimated in each group. Secondly, we explored these structural covariances from the whole brain level and brain ROI level. For the whole brain level, whole brain variability properties were estimated across all age groups. We also performed over-representation of correlation analysis (ORCA). Additionally, we compared the covariance of first group with those in other groups. For brain ROI level, we calculated the associations between structural covariance, age, cognition, and longevity-PRS, obtaining three association matrices.

### 3.2 Whole brain variability properties of structural covariance across age groups

For the cortical thickness covariance, the variance of whole brain pair-wise correlations increased significantly with age (r = 0.56, p = 2.17e-10), and the proportion of pairs that differed significantly also increased with age (r = 0.40, p = 1.65e-04). The entropy decreased significantly with age (r = −0.67, p = 2.86e-12). Age related decrease in the entropy implied overall reduction in the correlation between ROIs. The subcortical covariance showed similar trends in comparison with those in cortical thickness (Figure 2, Table S3).

**Figure 2.**
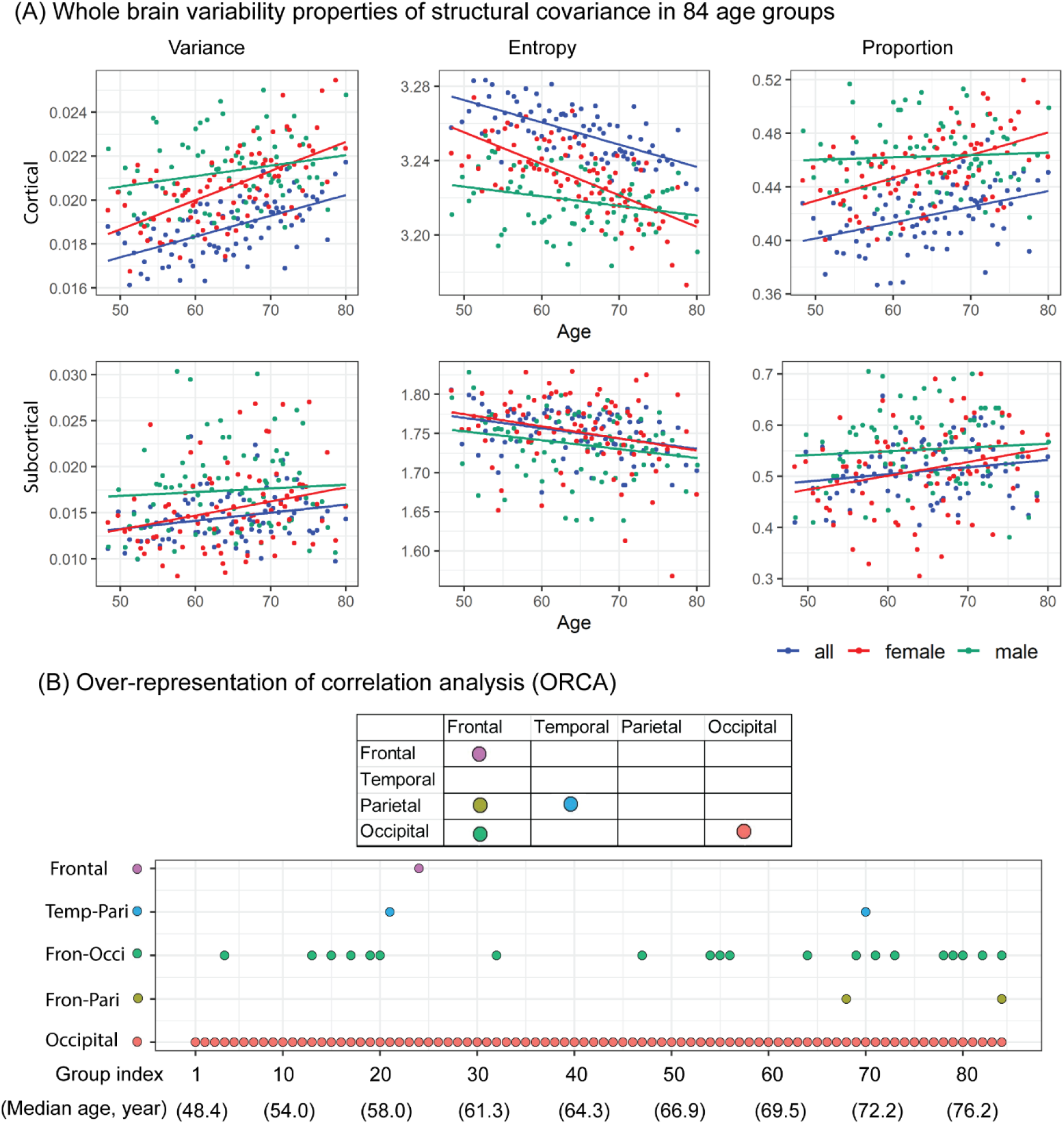
Whole brain variability properties of structural covariance and over-representation of correlation analysis (ORCA). (A) Whole brain variability properties of structural covariance in 84 age groups. Three global measures: variance, entropy, and proportions of pairs that differ significantly were estimated both in cortical thickness covariance and subcortical volume covariance. The points represent the corresponding measure in each age-group. Here, male, female, and the combined participants were investigated separately. (B) Over-representation of correlation analysis (ORCA). The x-axis represents 84 age groups, and the dots located in different positions indicate that significant enrichment of correlation coefficients between or within lobes were found in the specific age groups. Note that more correlations existed within occipital lobe than expected by chance, and this phenomenon was stable across all age groups. Significant enrichment of pairwise correlations was also observed between brain regions in frontal and occipital lobe.

We applied independent samples t-test to estimate the sex differences in those brain variability properties. Male and female participants showed significant differences after false discovery rate (FDR) correction (Benjamini and Hochberg, 1995) in cortical variance (p = 0.007), entropy (p = 5.89e-05), and the proportion of pairs that differ significantly (p = 0.031), as well as subcortical variance (p = 0.007), entropy (p = 0.025), and the proportion of pairs that differ significantly (p = 0.004). Cortical variances of females started lower than those of males for the younger age groups of our sample, then increased with age with a greater gradient than that of males, and finally surpassed the variances of males at around the age of 70. The faster rates of entropy and proportion of pairs that differ significantly in relation to increasing age in the female were also observed (Figure 2).

### 3.3 Test for ORCA

In general, ROIs within occipital lobe showed higher correlations across the all the age groups (for example, Figure S1). This is also statistically significant as confirmed by the ORCA analysis. The occipital lobe had significant enrichment of correlation coefficients above a threshold than by chance throughout all the ages (adjusted p < 0.05). Also, there were a greater number of significant negative correlations between frontal and occipital lobe (Figure S1). Hence ORCA showed significant enrichment for higher correlation (negative) between occipital and frontal lobes in some age groups, especially at older age groups (Figure 2B). The detailed statistics including the p-values can be found in Table S4.

### 3.4 Test for equality of correlation coefficients between the youngest group and other groups

By comparing the elements of correlation matrix in the first age group with corresponding elements of the correlation matrices of other age groups, we found that the correlation matrices of cortical thickness in older age were more significantly different from that in the youngest group, especially after the age of around 64, with females showing more pronounced trends than males in both cortical thickness and subcortical volume matrices (Figure 3). Additionally, by comparing the elements of correlation matrix of males and females in each age group, we did not find any significant trend across all 84 age groups. The detailed statistics including the p-values can be found in Table S5.

**Figure 3.**
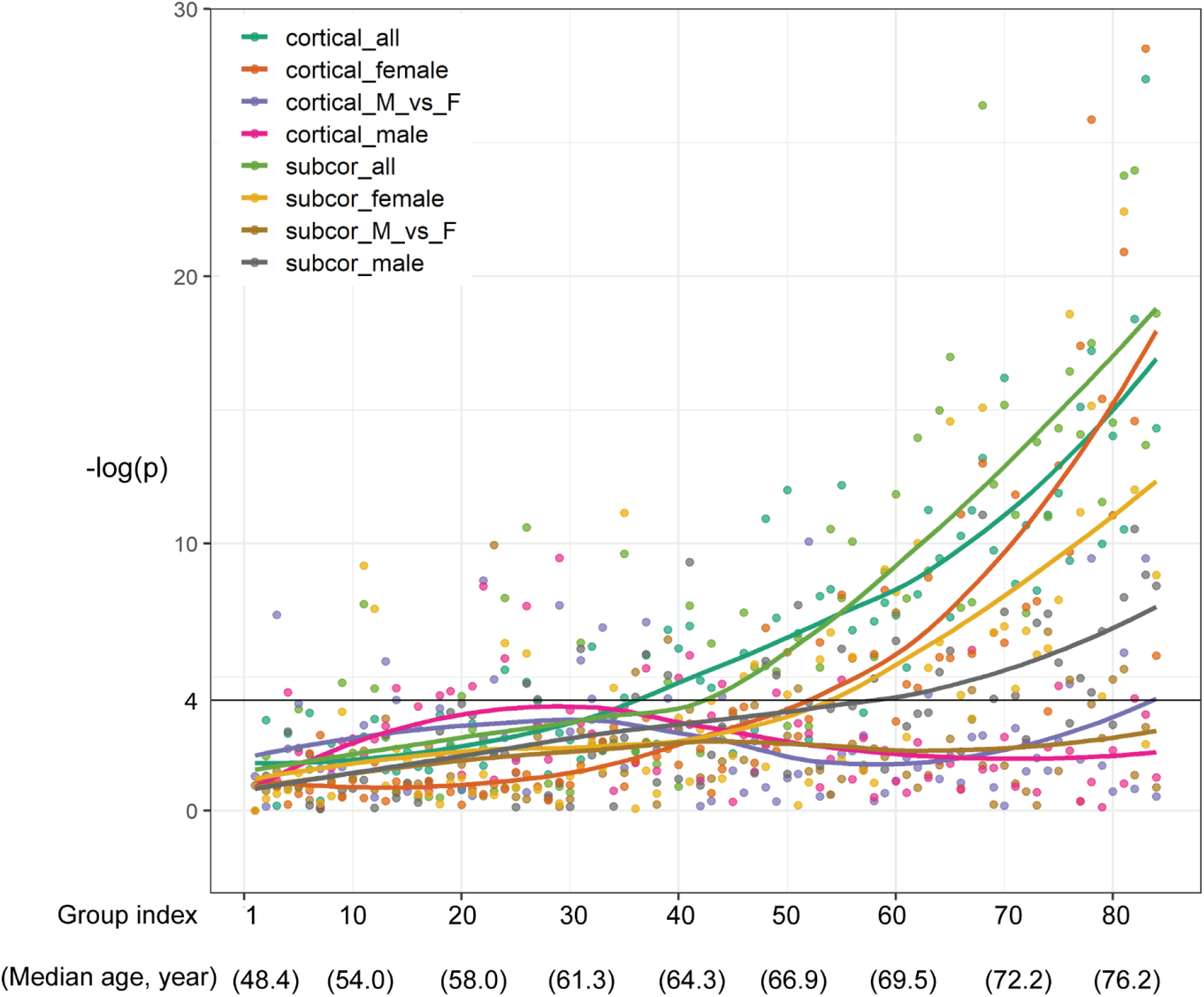
Test for equality of correlation coefficients between group 1 and each of other groups. Test for equality of correlation coefficients between group 1 and each of other groups. The x-axis indicates group index (from 1 to 84) and corresponding median age of each group. The y-axis indicates –log10(p) for the test for equality of group 1 matrix and each of other matrices. The black horizontal line (–log10[p-value] = 4.13) represents significant level p = 0.05 / (84*8) = 7.44e-05 after Bonferroni correction. The *cortical_all* represents the comparison of all participants’ correlation matrices between the first age group and each of other age groups (comparing group 1 with other groups). The *cortical_female* represents the comparison of only females’ correlation matrices between the first age group and each of other age groups (comparing group 1 with other groups). The *cortical_vs_F* represents the comparison of correlation matrices between males and females of the same age group (comparing males with females in each group). The *cortical_male* represents the comparison of only males’ correlation matrices between the first age group and each of other age groups (comparing group 1 with other groups). Other subcortical labels are similar to cortical labels.

### 3.5 Associations between structural covariance, age, cognition, and longevity-PRS

#### 3.5.1 Associations between structural covariance and age

##### Cortical thickness covariance and age

Each of the correlation coefficient across the age groups were tested for its association with median age. Sixty-two out of a total of 528 pairwise correlations were significantly associated with age across all 84 age groups after multiple test correction (Bonferroni correction, p<0.05/528 = 9.47e-5) (Figure 4-A). With an increasing age, some strongly negative correlations were observed between frontal lobe regions and other regions. The first four most significant pairwise correlations were found to be between transverse temporal and pars triangularis (r= 0.71, p = 3.92e-14), pars triangularis and superior frontal (r = −0.69, p = 2.31e-13), pericalcarine and superior frontal (r = −0.69, p = 3.06e-13), and between transverse temporal and superior frontal (r = 0.69, p = 6.13e-13) (Figure S2). The pairwise correlations between the regions within occipital lobe were all increased with an increasing age.

**Figure 4.**
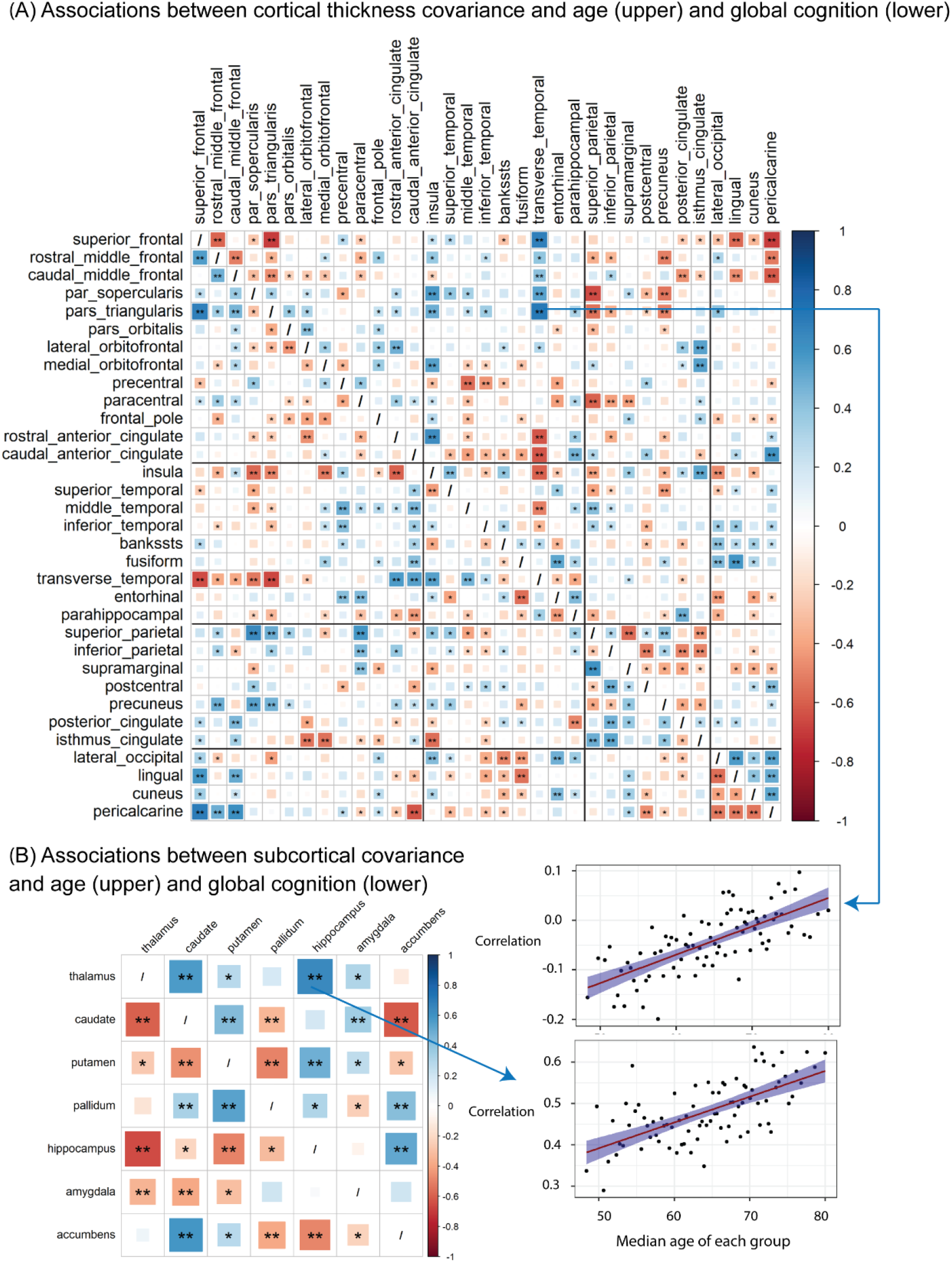
Associations between structural covariance and age/global cognition. Associations between structural covariance and age/global cognition in cortical thickness (A) and (B) subcortical volume. Covariance-age associations are shown in the upper right triangle and covariance-global cognition associations are shown in the lower left triangle. In the upper right triangle, every element in the matrix indicates the association between pairwise correlation of brain structures and age. In the lower left triangle, every element in the matrix indicates association between pairwise correlation of brain structures and global cognition. The single asterisk (*) represents the level of statistical significance p<0.05. Double asterisks (**) represents the associations remain statistically significant after Bonferroni correction.

##### Subcortical covariance and age

Ten out of a total of 21 pairwise correlations were significantly associated with age across all 84 age groups after Bonferroni correction (p<0.05/21=0.0024) (Figure 4-B). Notably, the correlations between hippocampus and other subcortical regions such as thalamus (r= 0.64, p = 3.57e-11), putamen (r= 0.49, p = 2.53e-06), and accumbens (r= 0.52, p = 4.43e-07) were significantly increased with increasing age. On the contrary, the correlation between caudate and accumbens was significantly decreased with increasing age (r = −0.62, p = 4.72e-10).

#### 3.5.2 Associations between structural covariance and cognition

##### Cortical thickness covariance and global cognition

The associations between pairwise correlations and global cognition were found to be largely opposite to their associations with age (Figure 4-A), i.e., greater pairwise correlations were associated with both older age and declined cognition. This can be explained by the fact that cognition usually declines in the ageing process (Supplementary, Figure S2). For example, an increased pairwise correlation between transverse temporal and pars triangularis was associated with increasing age (r= 0.71, p = 3.92e-14), but this correlation declined as global cognition increased (r = −0.66, p =6.07e-12). In addition, all the pairwise correlations within occipital lobe were significantly increased with older age and cognition decline. Figure S3 depicts the four most significant associations. The associations between covariance and cognitive domains (processing speed, executive function, and memory) were shown in Figures S4 to Figure S6. They all had similar association patterns.

##### Subcortical covariance and global cognition

Similar findings were seen in subcortical covariance (Figure 4-B). For example, the correlation between caudate and thalamus was positively associated with age (r = 0.57, p = 1.89e-08) but negatively associated with global cognition (r = −0.59, p = 2.57e-09). The associations between subcortical covariance and cognitive domains can be found in Figures S4 to Figure S6.

#### 3.5.3 Associations between structural covariance and longevity-PRS

We found a moderate association between pairwise brain region correlations and longevity-PRS. Nineteen out of a total of 528 pairwise correlations were significantly associated with longevity-PRS across all 84 age groups after Bonferroni correction (Figure 5). The pattern of associations between pairwise correlations and longevity-PRS were similar to the pattern of associations between pairwise correlations with age, partly because of the high positive correlation between longevity-PRS with age (Supplementary, Figure S2). For cortical thickness covariance, the correlation between pericalcarine and caudal middle frontal had the most significant association with longevity-PRS (r = −0.52, p = 3.83e-07). For subcortical covariance, the correlation between accumbens and caudate showed the most significant association with longevity-PRS (r = −0.42, p = 8.07e-05).

**Figure 5.**
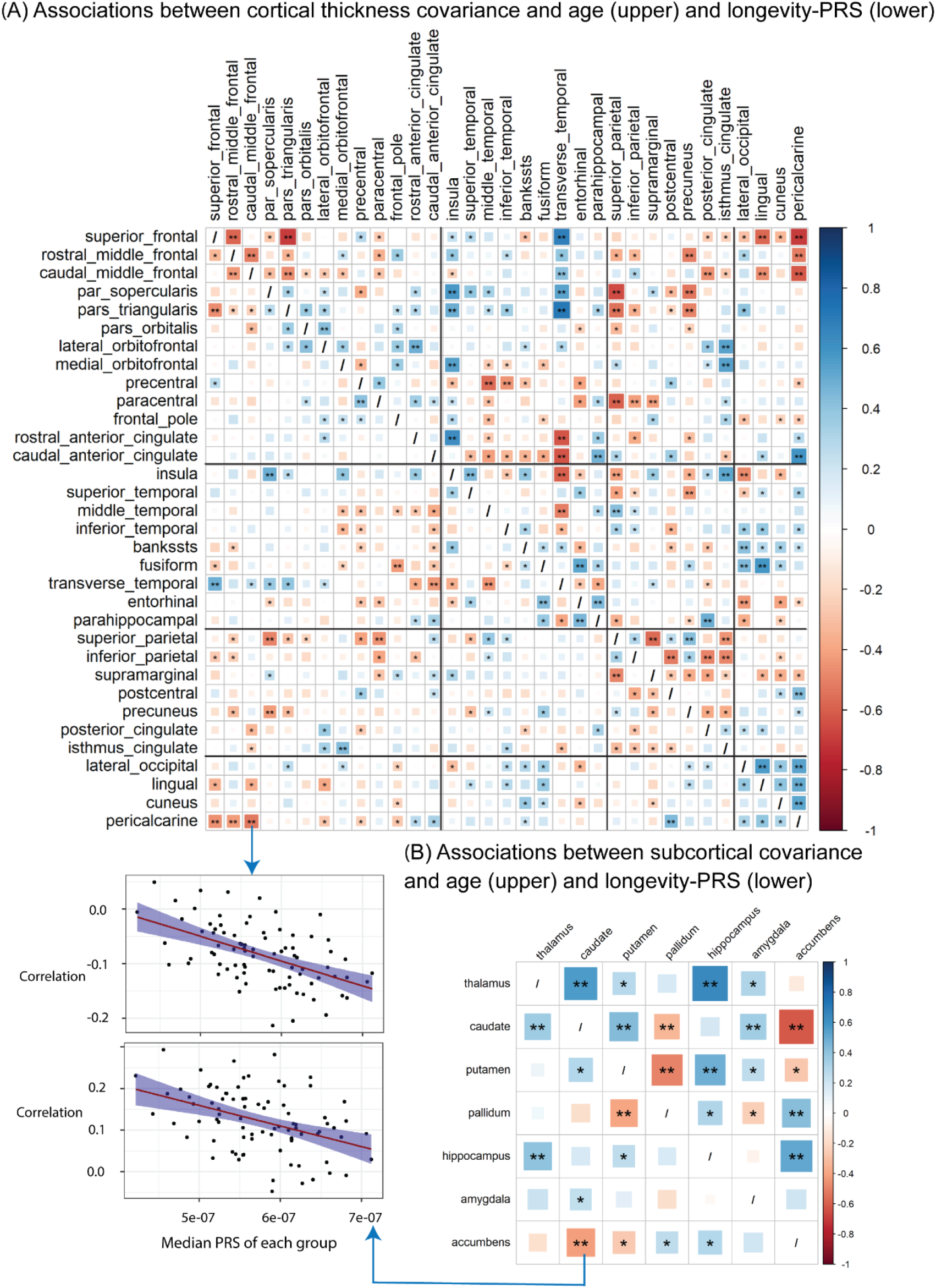
Associations between structural covariance and age/longevity-PRS. Associations between structural covariance and age/longevity-PRS in cortical thickness (A) and (B) subcortical volume. Covariance-age associations are shown in the upper right triangle and covariance-longevity-PRS associations are shown in the lower left triangle. The upper right triangle is the same as that in Figure 4. In the lower left triangle, every element in the matrix indicates association between pairwise correlation of brain structures and longevity-PRS. The single asterisk (*) represents the level of statistical significance p<0.05. Double asterisks (**) represents the associations remain statistically significant after Bonferroni correction.

The details about associations between pairwise brain region correlations and age, global cognition, and longevity-PRS can be found in Table S6 to Table S11.

## 4. Discussion

We investigated age-related structural covariance properties and their associations with cognition and longevity-PRS using a large sample of 42,075 participants drawn from the UK Biobank. First, the variability of whole brain structural covariance increased in older age. Second, there was significant and stable enrichment of pairwise correlations within the regions of the occipital lobe in the ageing process. Third, both cortical thickness and subcortical volume covariances in older groups were significantly different from those in the youngest group, especially for females. Fourth, brain structural covariances were not stable though the ageing process, with the pairwise correlations between some brain regions strengthening and some weakening with negative correlations in the relation to older age. Fifth, the significant pairwise correlations between brain regions were also significantly associated with cognitive ability, and weakly associated with longevity-PRS.

Our findings suggested that brain structural covariances had greater variability in older age, which was an indication of the loosening of cortical integration in the older brain. Given the interplay of different brain regions, which is similar to the changes of structural covariance in relation to the developmental stages in the childhood and adolescence (Váša et al., 2017; Vijayakumar et al., 2021), we hypothesised that the structural covariance in the ageing brain would also be ‘dynamic’ rather than ‘static’; and our results confirmed that. Additionally, the sex-related differences could be perhaps explained by the sex-related differences in brain function (Canli et al., 2002) and hormone regulation (Cosgrove et al., 2007). Overall, our findings suggested that the cortical integration appears to weaken in ageing process.

By comparing the corresponding element between the covariance matrix of the youngest group and the covariance matrix of an older group for all the older groups, it was found that the older the age group was, the more significant the difference it had in comparison with the youngest group. There was a noticeable acceleration of increase of the difference at around 64 (age group #40), which may indicate that the changes of human brain covariance start to accelerate at around the age of 64. This finding is in line with the findings of degenerative dementias such as Alzheimer’s disease whose rate of onset increases exponentially after the age of 65 (Fox and Schott, 2004). Additionally, the brain tissue loss has been reported to accelerate after that age (Resnick et al., 2003). It is worth noting that the structural covariances for females in older age diverged more from the youngest group than the covariances in males. This sex-related divergence was also evident in the differential rate of age-related increase of variance of whole brain covariances between males and females. Females started lower than the males but with a greater gradient. They then surpassed the males at the age of 70 (Figure 2). Diminishing estrogen in women after menopause was suggested (Morrison et al., 2006), as higher estradiol levels may protect the female brain against atrophy and white matter hyperintensity (Alqarni et al., 2021), and were also correlated with better structural network covariance and cognitive performance (Zsido et al., 2019).

The correlations between frontal lobe with other brain regions such as temporal, parietal, and occipital lobes changed significantly with age. For example, the correlations in superior and middle frontal-parietal, superior and middle frontal-occipital lobe were significantly decreased with increasing age. This suggested that cortical thickness in these regions was less correlated, and their interplay was steadily weakening in the ageing process. Similar decreased associations can be found between anterior cingulate and transverse temporal. On the other hand, during the ageing process, the transverse temporal regions showed increased correlation with superior and middle frontal regions, indicating that the transverse temporal region has different interactions with different frontal regions. The fact that the existence of both positive and negative correlations in relation to older age between certain brain regions suggested that there were concurrent strengthening and loosening of coordinated changes between these brain regions. Given that brain structures generally decline in the ageing brain, such as cortical thinning and subcortical atrophy, our finding demonstrated that there was a diverse range of rates of ageing in different brain structures. Notably, previous studies have shown that most of the above regions, for instance, regions in prefrontal, parietal lobe, and occipital, are highly correlated with each other within lobes and they are also genetically correlated with each other (Grasby et al., 2020; Hofer et al., 2020). We found that the regions in the occipital lobe were highly correlated with each other, and it has been reported that they share similar genetic mechanisms (Hofer et al., 2020). This could also perhaps explain our ORCA results of the enrichment of correlation coefficients within occipital lobe. Our findings indicated that although the enrichment in occipital correlations was significant though the ageing process, these correlations were significantly associated with age as well.

Like cortical thickness, subcortical volumetric covariances were also associated with the ageing process. As one of the important subcortical regions, the hippocampus plays a critical role in memory and learning, as well as spatial navigation (Burgess et al., 2002). It is vulnerable to neurodegenerative diseases, especially Alzheimer’s disease (Mu and Gage, 2011). In our analysis, the correlations between hippocampal volume with thalamus, putamen, and accumbens, were all significantly increased with increasing age, suggesting a synchronised decline in their volumes during the ageing process. The significantly decreased correlations between accumbens and caudate, putamen and pallidum, caudate and pallidum may indicate that these pairs of subcortical structures could have independent trajectories during ageing process. The amygdala and hippocampus are key components of the medial temporal lobe, and are involved in emotional perception and regulation (Groen et al., 2010). Our study showed that the covariance between these two regions remained relatively stable during ageing process.

We found significant associations between structural covariance and global cognition, processing speed, executive function, as well as memory. However, only a few significant associations between structural covariance and longevity-PRS were found in the current study. Previous work has shown that structural covariance of the default network was associated with cognitive ability (Spreng and Turner, 2013), and the synergy in the human brain which can quantify information integration may have evolved to support higher cognitive function (Luppi et al., 2022). In contrast to the positive associations between subcortical covariances and age, many subcortical covariances were found to be negatively associated with global cognition across all 84 age groups, especially the correlations between hippocampus and thalamus and those between caudate and thalamus. Although the exact biological basis of structural covariance has not been well understood, their associations with cognition could help provide clues.

Our study has some limitations. First, our data were cross-sectional in nature, which would not allow causal interpretation for the relationships of structural covariances and the relationships between structural covariances, cognition, and longevity-PRS. Second, in order to explore the associations between structural covariance and age, we computed brain structural covariance in the group with 500 participants, the grouping was empirically explored and decided. Finally, the generalisability of our results may not extend to other racial/ethnic groups, as we restricted our analyses to British ancestry.

## Conclusion

Our study revealed varying regional interactions related to brain morphology, with structural covariances showing greater variability in older age groups and associated with cognition and longevity-PRS. These findings imply that age, cognition and longevity-PRS might contribute to brain structural covariance organisation and provide novel insight into how brain regions interplay with each other during the ageing process.

## Conflict of Interest

None declared.

## Acknowledgements

This research has been conducted using the UK Biobank Resource under Application Number 37103. This research was undertaken with the assistance of resources and services from the National Computational Infrastructure (NCI) Australia, which is supported by the Australian Government. C.D. is supported by Scientia Scholarship of University of New South Wales, Sydney. J.J. is supported by John Holden Family Foundation. We also thank Angie Russell for her assistance in the preparation of the manuscript.

## Notes

### Competing Interest Statement

The authors have declared no competing interest.

