## Supplementary for "Brain structural covariances in the ageing brain in the UK Biobank": Supplementary Figures.docx

### Figure S1. The cortical thickness covariance (correlation matrix) in group 1, 2, 83, and 84.

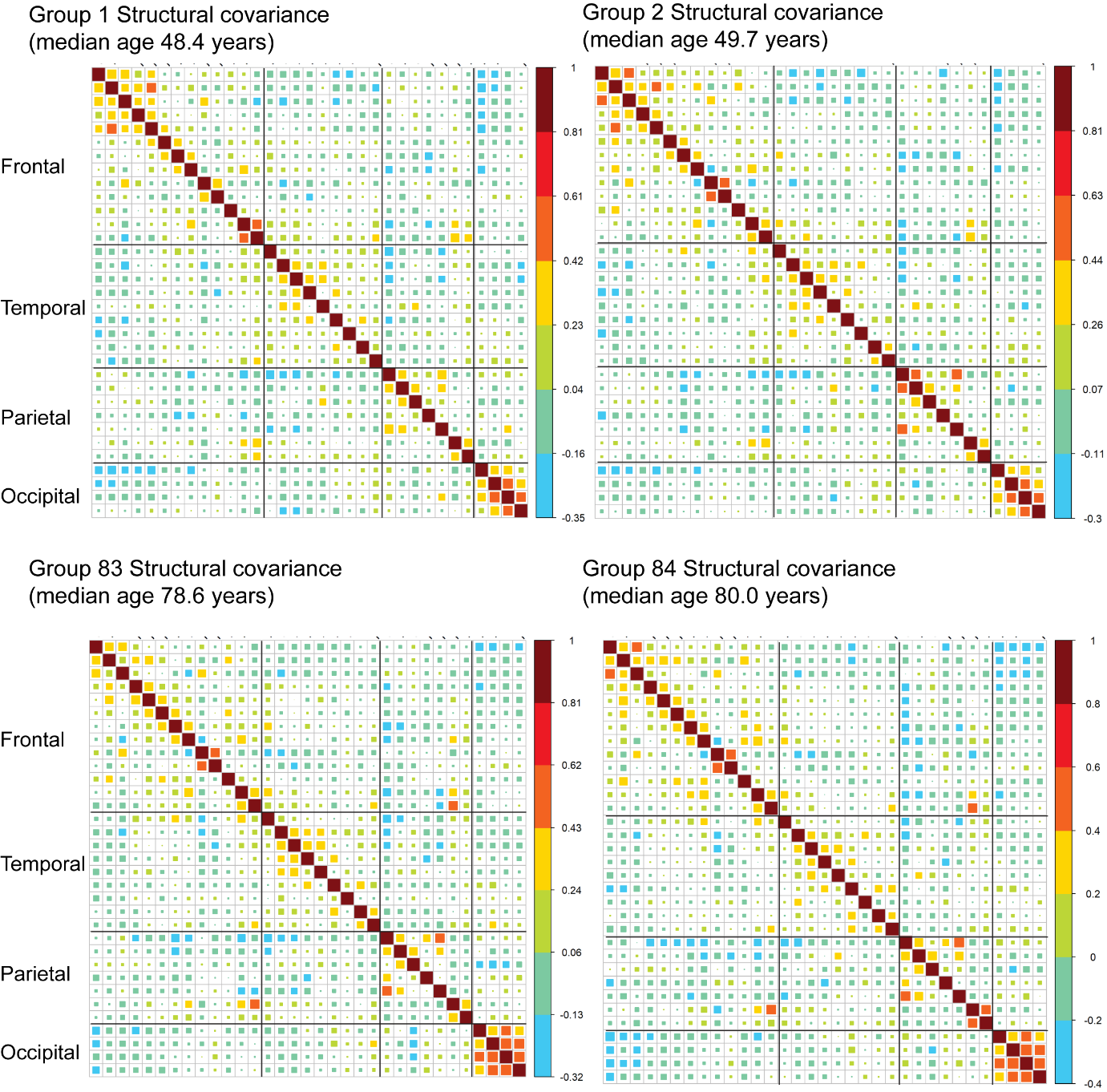

Figure S1. The cortical thickness covariance (correlation matrix) in group 1, 2, 83, and 84. The elements of each correlation matrix represent cortical thickness correlations between any two brain regions regressing out sex, scanner, and global mean cortical thickness.

### Figure S2. The association between global cognition and age, longevity-PRS and age across 84 age groups.

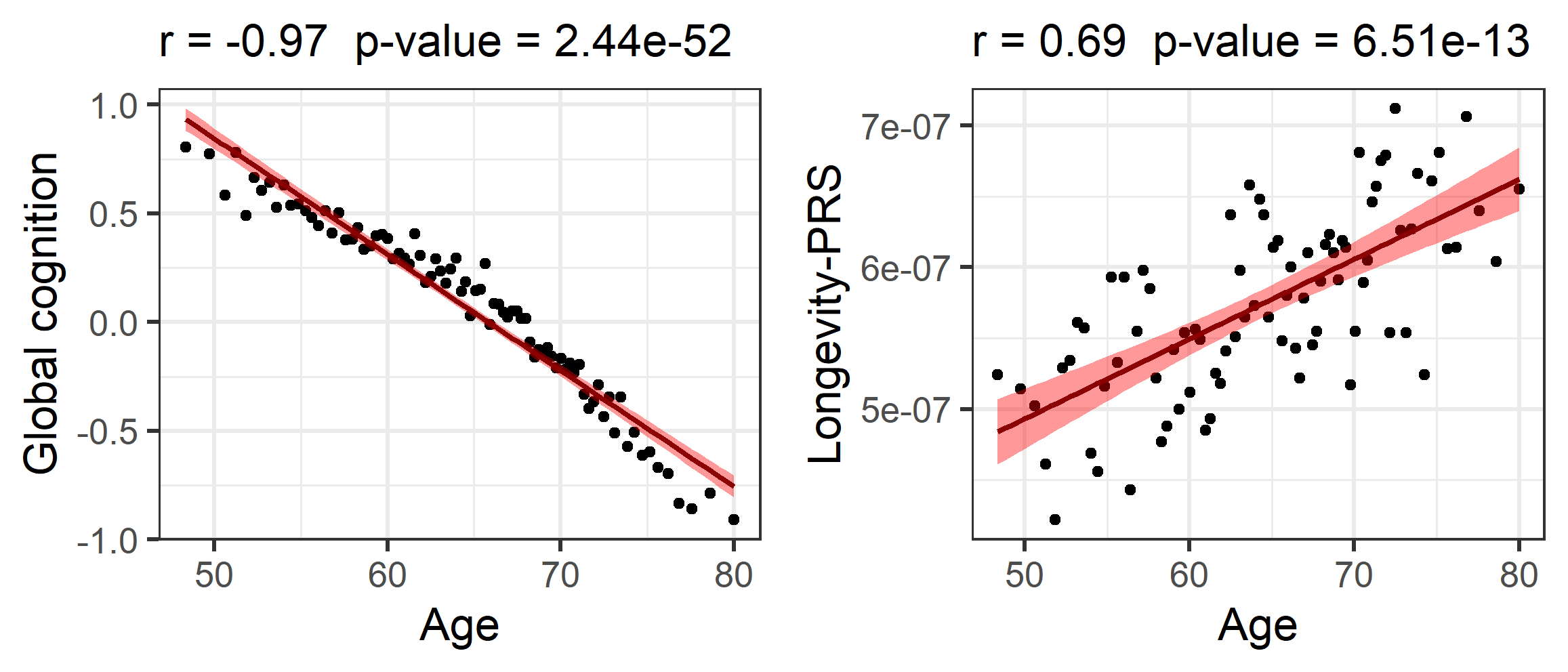

Figure S2. The association between global cognition (z-transformed) and age, longevity-PRS and age across 84 age groups. Age represents the median age in each age group. Similarly, global cognition and longevity-PRS represent the median values in each group.

### Figure S3. The first four associations between pairwise correlation and age/global cognition/longevity-PRS.

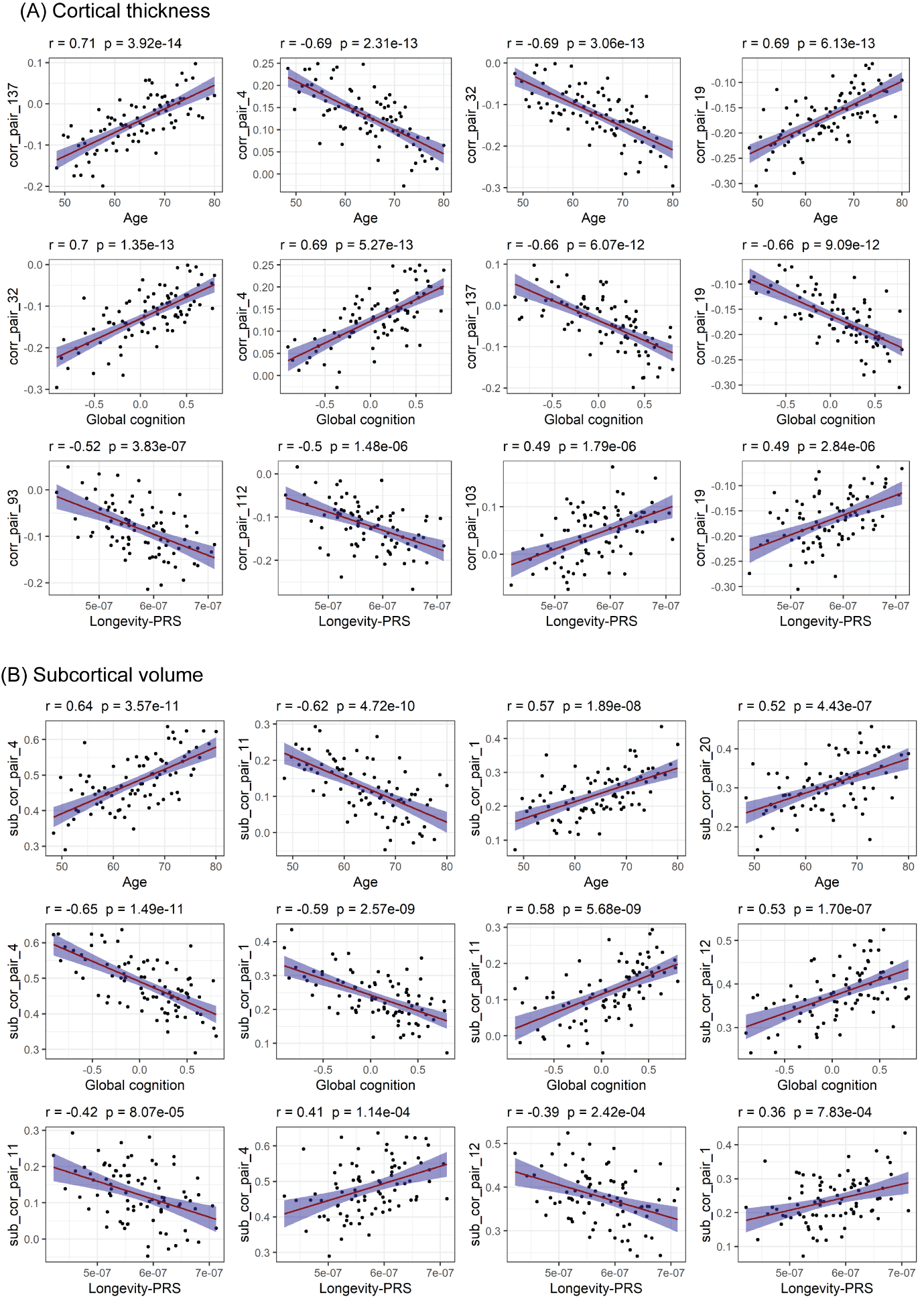

Figure S3. The first four associations between cortical pairwise correlation and age/global cognition/longevity-PRS. The x axis indicates median age/global cognition/longevity-PRS in each age group. The y axis indicates the cortical pairwise correlation, and corresponding index of pairwise correlation can be found in Table S1.

### Figure S4. Associations between structural covariance and age/processing speed.

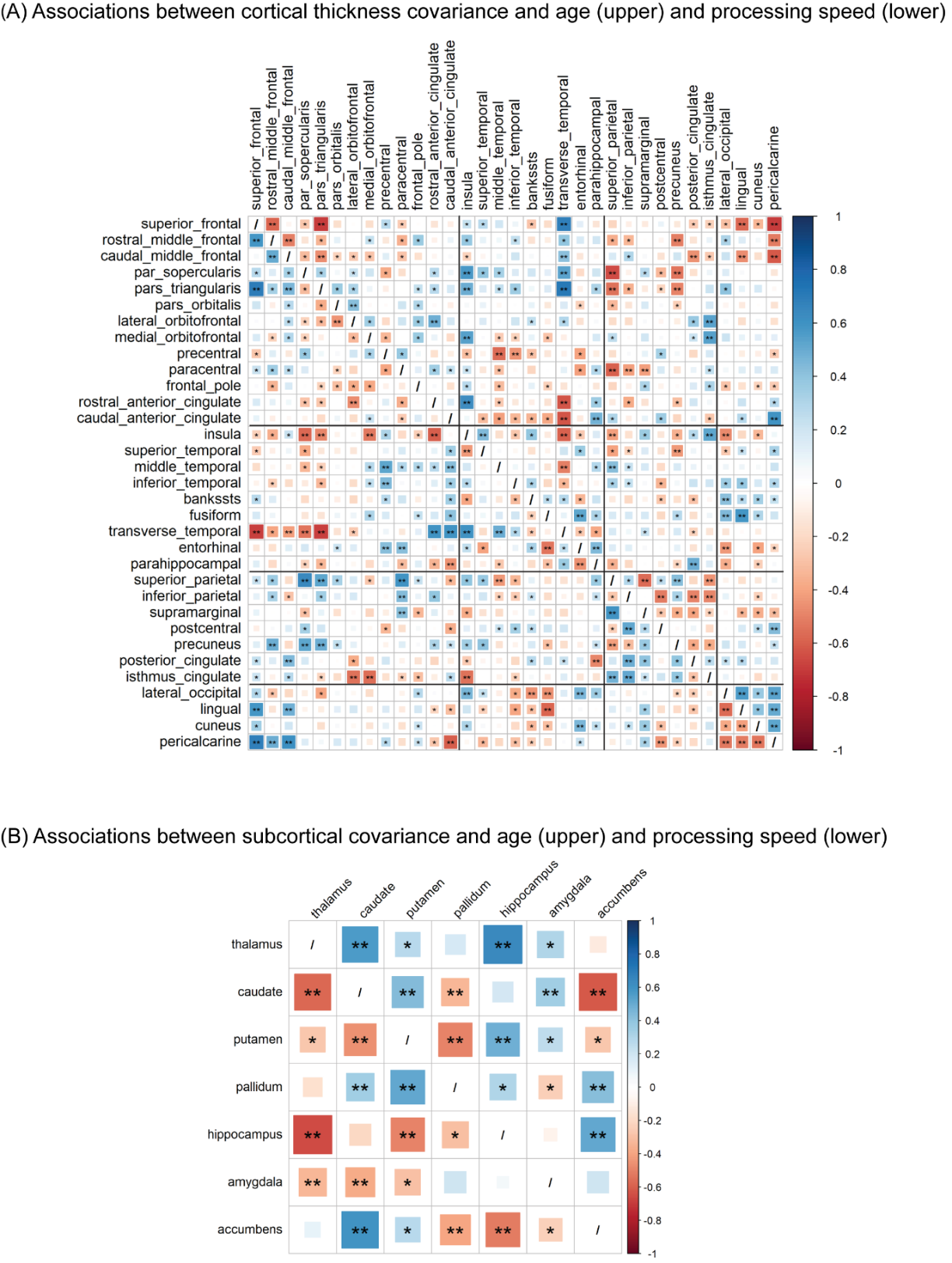

Figure S4. Associations between structural covariance and age/processing speed in cortical thickness (A) and (B) subcortical volume. Covariance-age associations are shown in the upper right triangle and covariance-processing speed associations are shown in the lower left triangle.

### Figure S5. Associations between structural covariance and age/executive function.

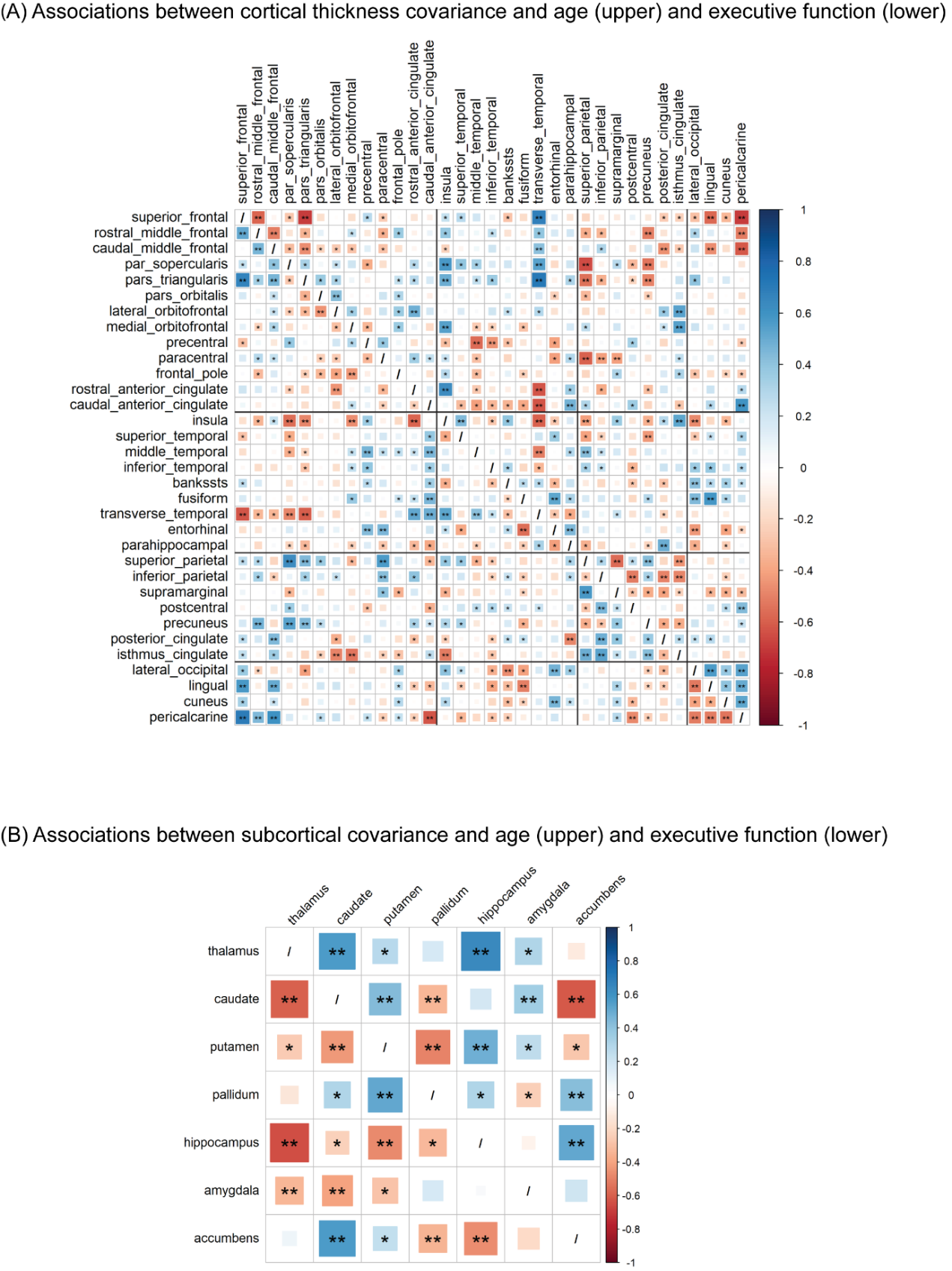

Figure S5. Associations between structural covariance and age/executive function in cortical thickness (A) and (B) subcortical volume. Covariance-age associations are shown in the upper right triangle and covariance-executive function associations are shown in the lower left triangle.

### Figure S6. Associations between structural covariance and age/memory.

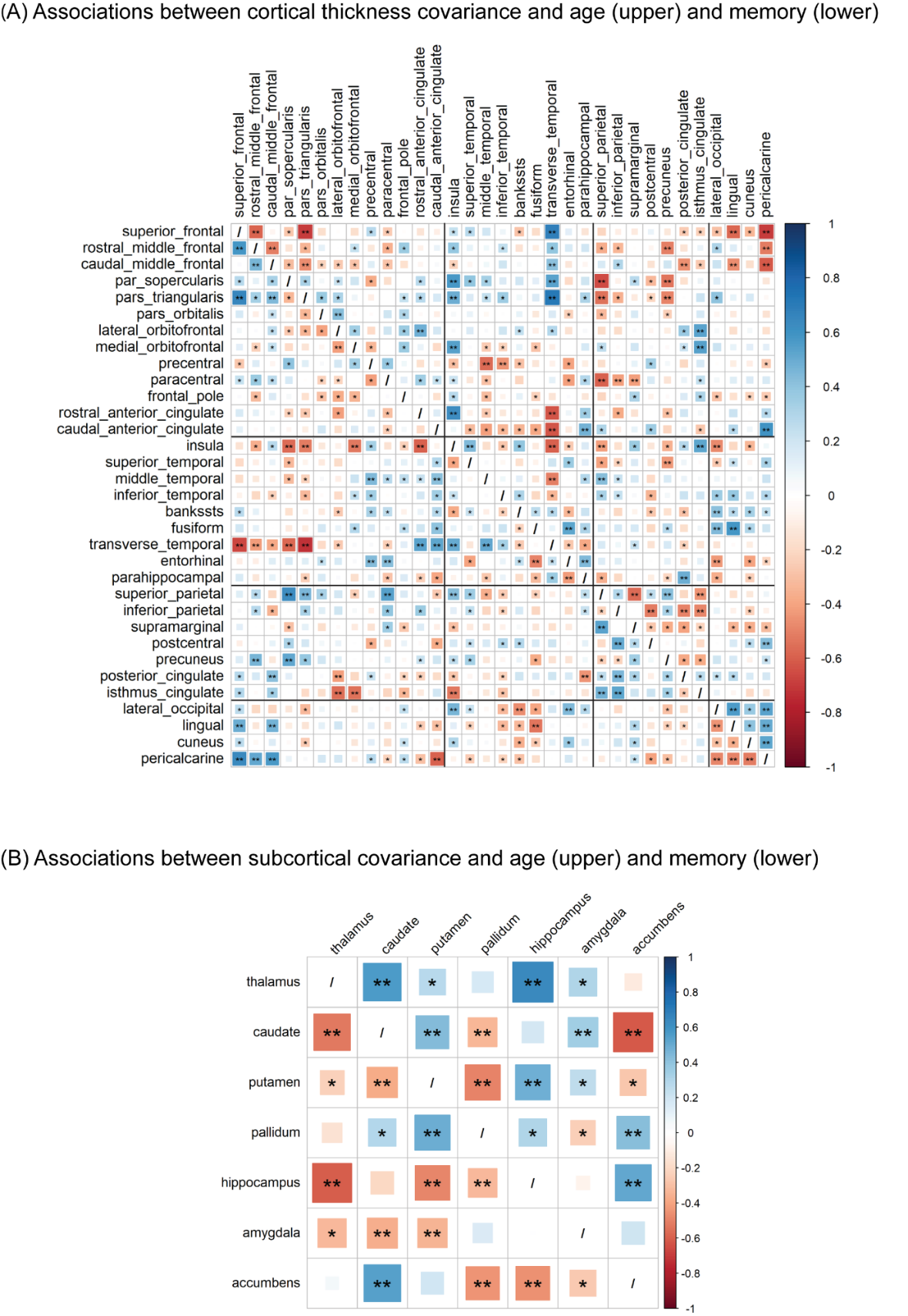

Figure S6. Associations between structural covariance and age/memory in cortical thickness (A) and (B) subcortical volume. Covariance-age associations are shown in the upper right triangle and covariance-memory associations are shown in the lower left triangle.
